# Wild-type allele of *TaHRC* suppresses calcium-mediated plant immune response by hijacking *TaCAXIP4* to trigger FHB susceptibility in wheat

**DOI:** 10.1101/2022.06.23.497398

**Authors:** Hui Chen, Zhenqi Su, Bin Tian, Guixia Hao, Harold N. Trick, Guihua Bai

**Affiliations:** Department of Agronomy, Kansas State University, Manhattan, KS 66506, USA; College of Agriculture and Biotechnology, China Agricultural University, Beijing 100193, China; Department of Plant Pathology, Kansas State University, Manhattan, KS 66506, USA; Mycotoxin Prevention and Applied Microbiology Research Unit, NCAUR, USDA-ARS, Peoria, IL 61604, USA; Hard Winter Wheat Genetics Research Unit, USDA-ARS, Manhattan, KS 66506, USA; Syngenta Crop Protection, Research Triangle Park, Durham, NC 27709, USA

**Keywords:** Wheat FHB, *Fhb1*, protein interaction, *TaHRC*, *TaCAXIP4*, Calcium-mediated plant immune response

## Abstract

Fusarium head blight (FHB) is one of the most destructive diseases of wheat worldwide. Epidemics of FHB cause a serious reduction in grain yield and quality of wheat and result in significant economic losses to wheat producers. Recently, we have cloned a histidine-rich calcium-binding protein gene (TaHRC) as the causal gene for Fhb1 and demonstrated that the wild-type allele of TaHRC conditions FHB susceptibility and a large deletion including the start codon resulted in FHB resistance. However, the molecular mechanisms on how TaHRC regulating FHB susceptibility remains unknown. In this study, we conducted yeast two-hybrid screening (Y2H) against the wheat cDNA expression libraries using TaHRC as bait and identified a cation exchanger (CAX)-interacting protein 4 (TaCAXIP4) as the candidate protein that interacts with TaHRC to affect calcium transport activity. The strong interaction was further confirmed by Bimolecular fluorescence complementation (BiFC) assays. Using gene editing, we edited three different sites (one before and one within and one after the NLS domain) of TaHRC in a susceptible wheat cultivar ‘Bobwhite’ using the CRISPR/Cas9 gene editing technology and demonstrated the N-terminus carrying NLS domain of TaHRC plays a critical role for the interaction and conditions TaHRC function on FHB susceptibility. We determined that the interaction between TaCAXIP4 and TaHRC occurs in the nuclei of cells by subcellular colocalization assay. Intriguingly, we found TaHRC can sequester TaCAXIP4 to suppress the Ca2+ transporting activity of TaCAX1 (a H+/Ca2+ antiporter) through yeast calcium suppression assay and suggested wild-type TaHRC may hijack TaCAXIP4 to suppresses calcium-mediated plant immune response during Fusarium infection in wheat. Furthermore, we performed the reactive oxygen species (ROS) assays and further showed that TaHRC might suppress the chitin-triggered plant immune responses during Fusarium infection by sequestering TaCAXIP4 to trigger FHB susceptibility, which facilitates the pathogen spread within a wheat spike. This work provides first line of evidence to support wild type Fhb1 is a susceptible gene and how Fhb1 wild type allele regulate FHB susceptibility.

---

Dear Editor,

Fusarium head blight (FHB) caused by *Fusarium graminearum* is a destructive disease in wheat (*Triticum aestivum*) worldwide. FHB epidemics significantly reduce wheat grain yield and quality and threat global food security and safety. Growing FHB resistant wheat cultivars can effectively minimize FHB damage (Bai et al., 2018). Among more than 600 quantitative trait loci reported for FHB resistance to date, only a few including *Fhb1* from Chinese sources show a major effect (Su et al., 2019; Zheng et al., 2020). More recently, we have cloned a histidine-rich calcium-binding protein gene (*TaHRC*) as the causal gene for *Fhb1* and demonstrated that the *TaHRC* wild-type allele conditions FHB susceptibility and a large deletion including the start codon resulted in FHB resistance (Su et al., 2019). However, the molecular mechanisms on how *TaHRC* regulating FHB susceptibility remains unknown.

To identify TaHRC interacting proteins, we conducted yeast two-hybrid screening (Y2H) against the wheat cDNA libraries using *TaHRC* as a bait. After screened 130 million clones, we identified 25 high confidence candidate interacting proteins (HCIP’s) (Supplemental Table S1). Since previous studies showed that HRC regulates Ca^2+^-uptake and -release to maintain Ca^2+^-homeostasis and a cation exchanger (CAX)-interacting protein 4 (TaCAXIP4) is the only one that associated with calcium transport activity among the 25 HCIP’s (Cheng et al., 2002), we selected TaCAXIP4 to investigate its interaction with TaHRC in yeast. We first cloned the full-length coding sequences (CDS) of *TaCXIP4* and *TaHRC* from ‘Clark’ (an *Fhb1* susceptible wheat cultivar) and then constructed the CDSs into a prey vector (pGADT7-TaCAXIP4 with a leucine report gene) and a bait vector (pGBKT7-TaHRC with a tryptophan report gene), respectively. We also constructed additional bait vectors for the *TaHRC* N-terminal fragment containing/without a nuclear localization signal (NLS) domain (pGBKT7-TaHRC-N and pGBKT7-TaHRC-NΔNLS) and the C-terminal fragment without NLS domain (pGBKT7-TaHRC-C) (Figure 1, a), and then co-transformed them into a Y2HGold yeast strain. All of the yeast cells grew well on the medium lacking Leucine and Tryptophan (Figure 1, b, left panel), indicating successful co-transformation. However, only the yeast cells co-expressing the full-length or N-terminus of *TaHRC* with *TaCXIP4* grew on the selective medium (Figure 1, b, right panel), suggesting a strong interaction between the two proteins in yeast and that the *TaHRC* N-terminus carrying NLS domain is essential for the interaction.

**Figure 1.**
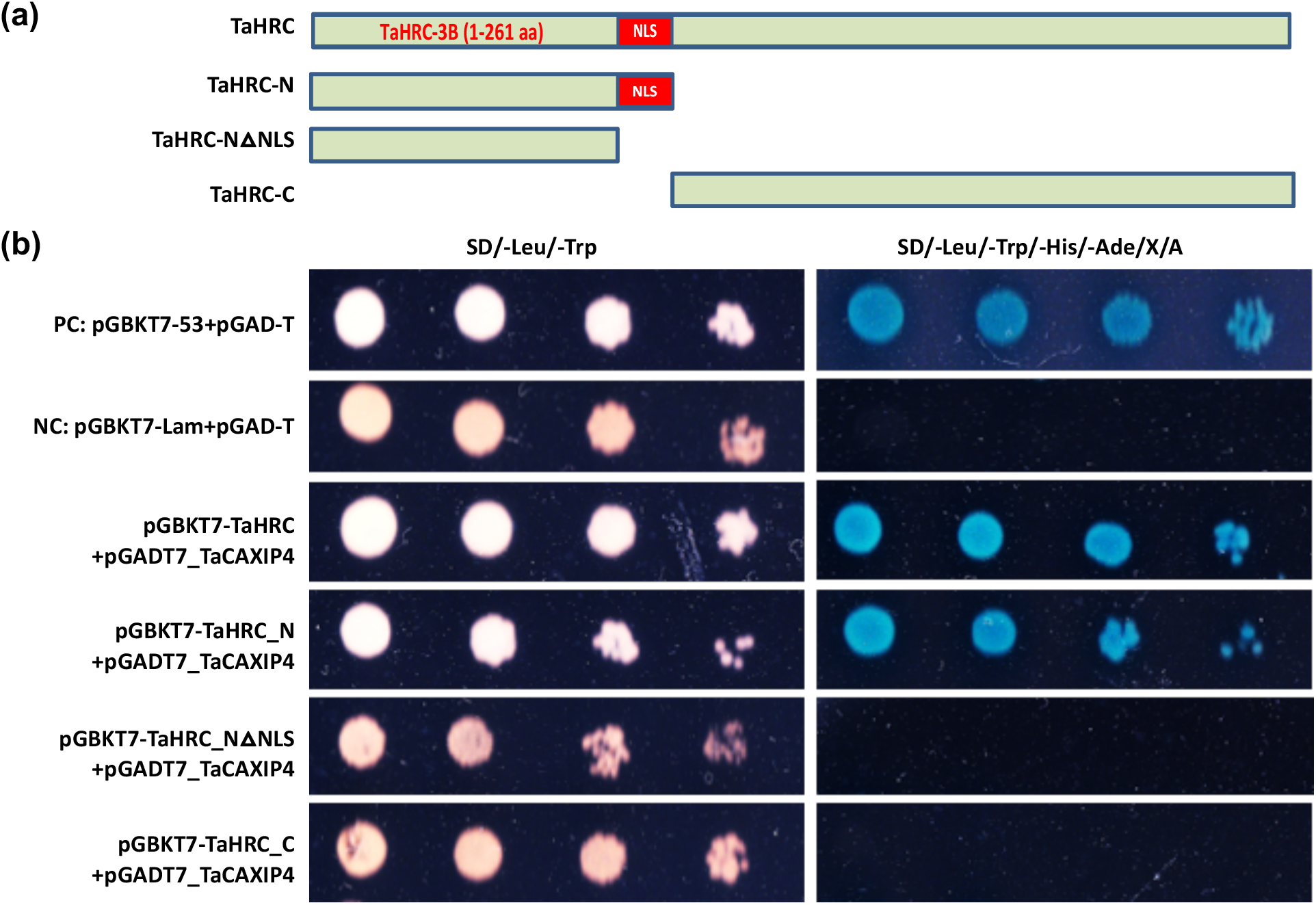
Confirmation of the interaction between TaHRC and TaCAXIP4 in yeast. a, Schematic presentation of different *TaHRC* constructs used for yeast transformation. b, Co-transformation assays to validate the TaHRC-TaCAXIP4 interaction in yeast. Five microliters of serial dilutions of yeast cells were spotted onto the synthetic dextrose (SD) medium. SD/-Leu/-Trp indicates SD medium lacking leucine (Leu) and tryptophan (Trp). SD/-Leu/-Trp/-His/-Ade/X/A indicates SD medium lacking Leu, Trp, adenine (Ade) and histidine (His), but contains 20 ng/μL of X-alpha-Gal and 125 ng/ml of Aureobasidin A (AbA). PC is the positive control by co-transformation of pGADT7 and pGBKT7-53; NC is the negative control by co-transformation of pGADT7 and pGBKT7-lam.

To confirm the critical role of N-terminus carrying NLS domain of *TaHRC* on FHB susceptibility, we edited three different sites (one before and one within and one after the NLS domain) of *TaHRC* in a susceptible wheat cultivar ‘Bobwhite’ using the CRISPR/Cas9 gene editing technology (Chen et al., 2022) and identified one mutant each at the three different target sites, respectively, with two insertion mutations and one deletion mutation (Figure 2, a). The homozygous M2 plants were inoculated with a conidiaspore suspension of *F. graminearum* (GZ3639) by single spikelet injection at early anthesis in a growth chamber (Su et al., 2019). The percentage of symptomatic spikelets (PSS) in a spike in the Mut01 and Mut02 with disrupted N-terminus carrying NLS domain was significantly reduced at 14 days after inoculation, whereas PSS of the Mut03 with the complete N-terminus carrying NLS domain did not change (Figure 2, b and c), suggesting that reduced FHB susceptibility in the mutants Mut01 and Mut02 with the disrupted N-terminus is likely due to abolished *TaHRC* function and the NLS domain in the N-terminus is critical for the interaction.

**Figure 2.**
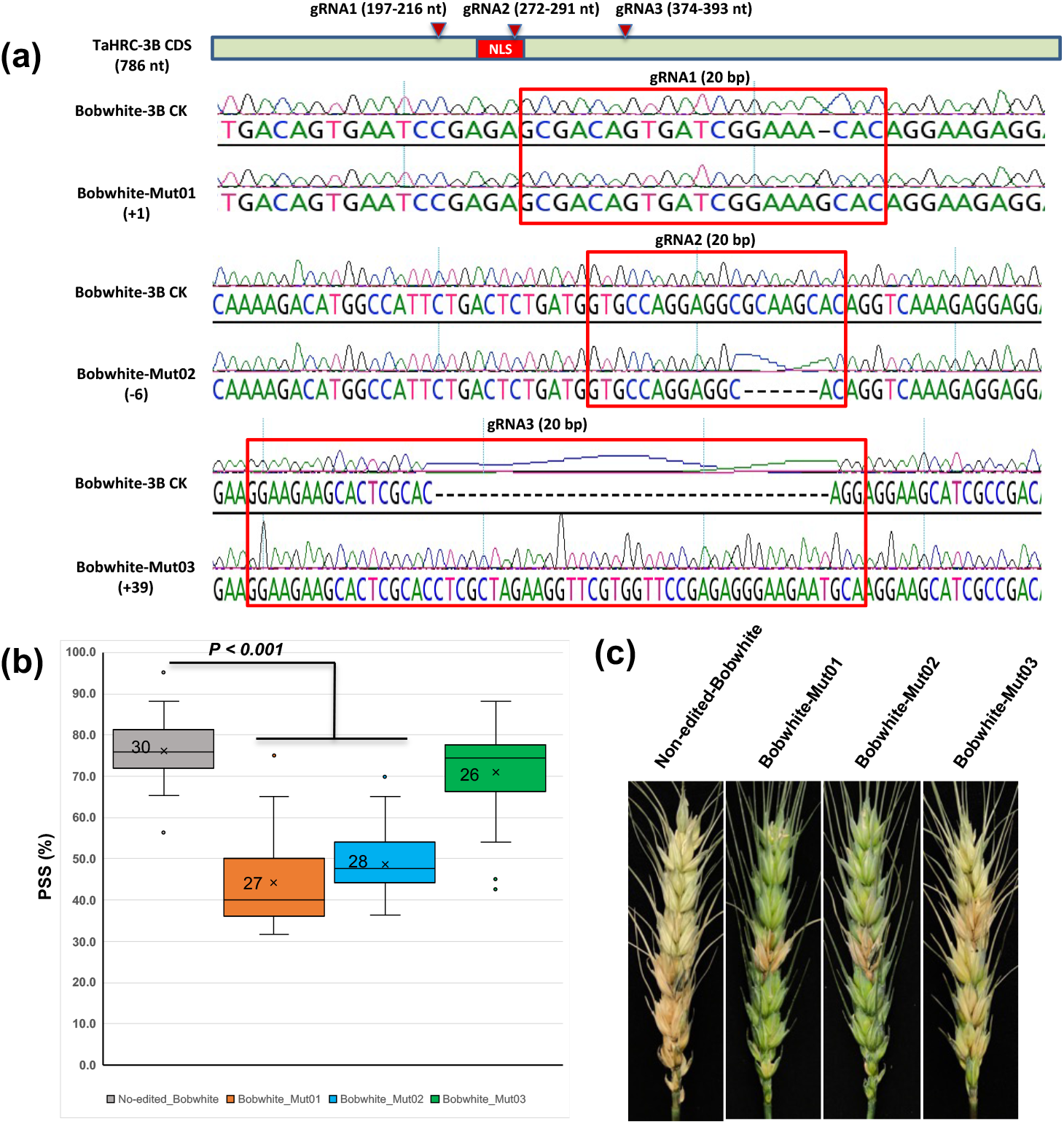
Determination of the functional role of the N-terminus carrying the NLS domain of TaHRC on FHB susceptibility using the CRISPR/Cas9 gene editing technology. a, Edited sequences at three different target sites of *TaHRC* as identified from the three Bobwhite mutant lines by Sanger sequencing. Bobwhite-Mut01 has one nucleotide insertion (+1), Bobwhite-Mut02 has six nucleotide deletion (−6) and Bobwhite-Mut03 has 39 nucleotide insertion (+39). The red bar within the coding sequences (CDS) of *TaHRC-3B* is a nuclear localization domain. The red arrows point to the targeted sites of three different gRNAs. b, Comparison of mean percentages of FHB symptomatic spikelets (PSS) between the three mutant lines and non-edited control plants. Boxes indicate the 25th–75th percentile, whiskers indicate the full data range, center lines indicate medians, crosses indicate means and the numbers inside boxes indicate sample size. *P*-values were generated from two-sided unpaired Student’s t-tests of the mean PSS of the mutant lines versus the mean PSS of the non-edited line. c, FHB symptoms in the inoculated spikes from the three mutant lines and non-edited Bobwhite control plants.

To validate the interaction between *TaHRC* and *TaCXIP4* in planta, we fused the full-length CDS of *TaHRC* and *TaCAXIP4* into the N-terminus and C-terminus of a split yellow fluorescent protein (YFP) in the expression vectors as YN-TaHRC and YC-TaCAXIP4, respectively (Lu et al., 2010), and infiltrated the mixed *Agrobacterium tumefaciens* cultures harboring equal concentrations of YN-TaHRC and YC-TaCAXIP4 vectors into 6-week-old epidermal *Nicotiana benthamiana* leaves for bimolecular fluorescence complementation (BiFC) assays. We observed strong signals of the reconstituted YFP fluorescence in the nuclei of cells at 48 hours after infiltration (Figure 3, a), indicating a very strong interaction between TaHRC and TaCXIP4 in planta.

**Figure 3.**
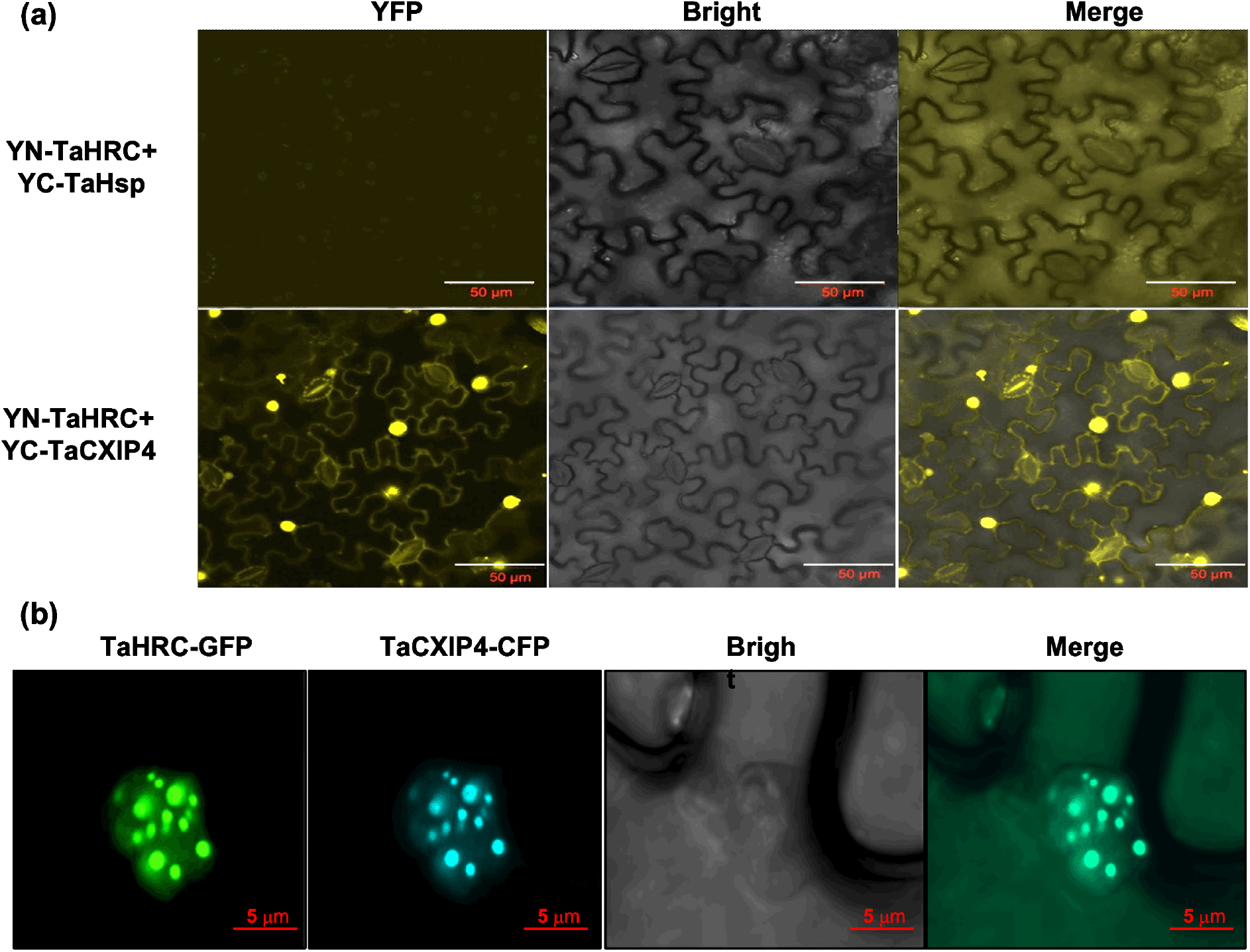
Confirmation of the interaction between TaHRC and TaCAXIP4 in planta. a, Bimolecular fluorescence complementation (BiFC) assays to show the protein interaction between TaHRC and TaCAXIP4 in planta. The reconstituted YFP signals were observed under a Zeiss LSM 880 confocal microscope (Carl Zeiss, Germany). An unrelated heat shock protein of YFP fusion (YC-TaHsp) was used as the negative control. Scale bars = 50 μm. b, Subcellular colocalization of TaHRC and TaCXIP4 in the nuclei of plant cells. After infiltration with the mixed *Agrobacterium* cultures harboring equal concentrations of TaHRC-GFP and TaCAXIP4-CFP vectors into 6-week-old epidermal *N. benthamiana* leaves at 48 hours, the fluorescence signal of GFP and CFP were imaged under a Zeiss LSM 880 confocal microscope in two channels and merged using a lookup table with raw data in green and cyan colors, respectively. Scale bars = 5 μm.

To determine where the interaction occurs in plant cells, we fused the CDS of *TaHRC* and *TaCAXIP4* into the N-terminus of an intact green fluorescent protein (GFP) and a cyan fluorescent protein (CFP) in the expression vectors as TaHRC-GFP and TaCXIP4-CFP, respectively, and infiltrated the mixed *A. tumefaciens* cultures harboring equal concentrations of TaHRC-GFP and TaCXIP4-CFP vectors into 6-week-old epidermal *N. benthamiana* leaves for subcellular colocalization assays. At 48 hours after infiltration, the strong GFP and CFP fluorescence signals from the co-expression of TaHRC-GFP and TaCXIP4-CFP fusion proteins were observed in the nucleus speckles (Figure 3, b), confirming their interaction in the nuclei.

The *Arabidopsis* cation exchanger 1 (CAX1) is a H^+^/Ca^2+^ antiporter that plays an important role in maintaining cellular Ca^2+^ homeostasis, whereas CAX1 activity is activated by CAXIP4 (Cheng et al., 2004). To investigate whether the HRC-CXIP4 interaction affects the ability of CXIP4 to activate CAX1 Ca^2+^ transport activity, we cloned the full-length CDS of *TaCAX1* and *TaCAXIP4* from Clark into the yeast expression vector pGBKT7 and transformed/cotransformed the constructs (pGBKT7-TaHRC, pGBKT7-TaCAXIP4 and pGBKT7-TaCAX1) into a Ca^2+^ sensitive yeast strain K667 (hypersensitive to high concentrations of Ca^2+^). Yeast cells expressing/coexpressing *TaHRC*, *TaCAXIP4* and *TaCAX1* were assayed on a yeast extract/peptone/dextrose (YPD) medium supplemented with and without 200 mM CaCl_2_, respectively. The K667 cells expressing *TaCXIP4* or *TaCAX1 or TaHRC* alone did not grow on the medium containing 200 mM CaCl_2_, whereas the K667 cells coexpressing both *TaCXIP4* and *TaCAX1* grew well on the same medium (Figure 4, a), indicating TaCXIP4 activated TaCAX1 to mediate the Ca^2+^ transport activity in yeast. However, the K667 cells coexpressing *TaCAX4, TaCAX1* and *TaHRC* did not grow on the medium with 200 mM CaCl_2_, indicating *TaHRC* sequestered *TaCAXIP4* to suppress the Ca^2+^ transporting activity of *TaCAX1*. These results suggested that wild-type *TaHRC* may hijack *TaCAXIP4* to suppresses calcium-mediated plant immune response during *Fusarium* infection in wheat.

**Figure 4.**
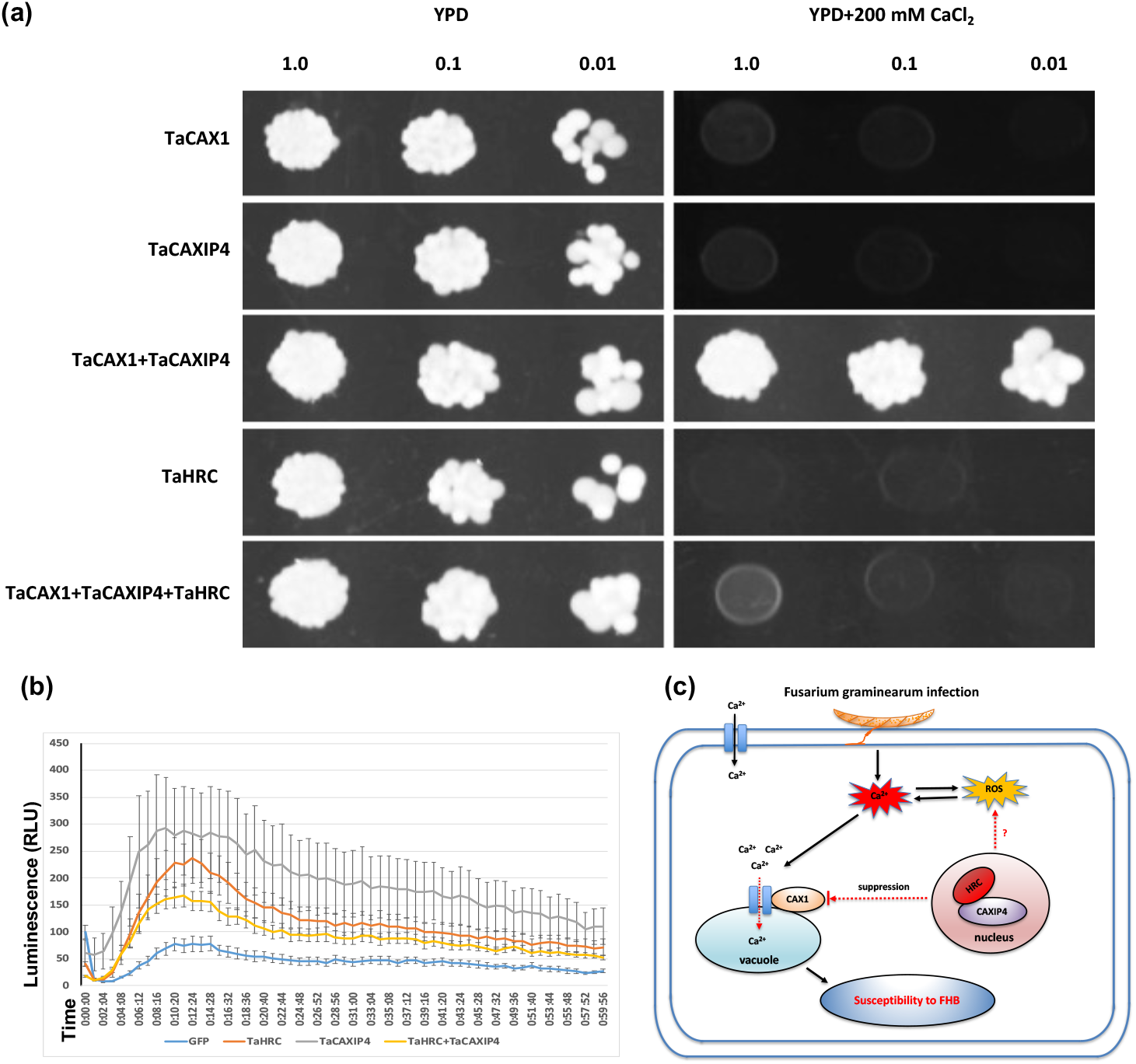
Effect of the HRC-CXIP4 interaction on CAX1-mediated Ca^2+^ transport activity and chitin-triggered plant immune responses during *Fusarium* infection. a, Suppression of K667 yeast Ca^2+^ sensitivity in cells coexpressing various combinations with three plasmids, *TaCAXIP*4, *TaCAX1* and *TaHRC* as indicated. Yeast cells were grown to A600 =1.0 in selection medium at 30 °C. Five microliters of serial dilutions were spotted onto YPD medium supplemented with/without 200 mM CaCl_2_. Photographs were taken after 3 days of culture. b, Chitin-induced reactive oxygen species assay via transient expression of *TaHRC-GFP* and *TaCAXIP4-GFP* in *Nicotiana benthamiana* leaves. Luminescence was measured in 200 μl of the assay solution (17 mM lumino, 1 μM horseradish peroxidase, and crab shell chitin at 200 μg/ml) for 60 min. Leaves expressing *GFP* served as the positive control and the assay solution without chitin served as the negative control. Lines are means and standard error with n = 12. Assays were repeated four times with similar results. RLU = relative light unit. c, A proposed model of the calcium-mediated plant immune response suppressed by wild-type allele of *TaHRC* to enhance wheat FHB susceptibility. *TaHRC* hijacks *TaCAXIP4* to suppress the Ca^2+^ transporting activity of *TaCAX1* and disrupte the Ca^2+^ signal transduction during the immune response to *F. graminearum* infection.

Production of reactive oxygen species (ROS) is critical for successful activation of plant immune responses against pathogens (Hao et al., 2019). To determine the role of *TaHRC* in regulating plant immunity, we infiltrated *N. benthamiana* leaves with *Agrobacterium* cultures harboring the full-length CDSs of *TaHRC-GFP* and *TaCAXIP4-GFP* in the expression vectors, and observed a high level of chitin-trigged ROS in the plants expressing *TaHRC* or *TaCAXIP4* alone, but a low level of ROS in the plants coexpressing both *TaCAXIP4-GFP* and *TaHRC-GFP* (Figure 4, b), which suggested that *TaHRC* might suppress the chitin-triggered plant immune responses during *Fusarium* infection by sequestering *TaCAXIP4* to trigger FHB susceptibility.

In summary, we demonstrated that *TaCAXIP4* interacts with *TaHRC* in the nuclei of cells to trigger wheat FHB susceptibility and the functional NLS domain in *TaHRC* N-terminus is essential for the interaction. *TaHRC* may hijack *TaCAXIP4* to suppresses calcium-mediated plant immune responses, which facilitates the pathogen spread within a wheat spike (Figure 4, c). This study provides further insights into molecular mechanisms of *TaHRC* on regulating FHB susceptibility in wheat.

## Data availability

Data that support the findings of this work are included in the parper and its Supplemental Information files.

## Acknowledgments

We would like to extend our thanks to Dr. Kendal D Hirschi and Dr. Jian Yang at Baylor College of Medicine for providing the yeast strain K667 and to the confocal core facility in the College of Veterinary Medicine at Kansas State University for the assistance and usage of the confocal microscopes.

## Author Contributions

H. C. and G.B. designed the project, H. C., Z. S., G. H., and B. T. performed the experiments. H. C., H. T. and G.B. wrote the manuscript and all authors read and approved the final version of the manuscript.

## Supplemental information

### Materials and methods

#### Yeast Two-Hybrid (Y2H) screening and co-transformaiton assays

Y2H screening experiment was performed with the ULTImate Y2H screen system by Hybrigenics Services (Paris, France http://www.hybrigenics-services.com). The full-length coding sequences (CDS) of *TaHRC* was amplified from ‘Clark’ (an *Fhb1* susceptible wheat cultivar) and then constructed into a bait vector (pGBKT7-TaHRC). The construct was verified by Sanger sequencing and used as a bait to screen a wheat cDNA library fused to Gal4. The full-length CDS of *TaCXIP4* from ‘Clark’ was amplified and constructed into a prey vector (pGADT7-TaCAXIP4), two N-terminal truncated fragments of *TaHRC* (one with a NLS domain, 1-96 aa; another without NLS domain, 1-81 aa) and the C-terminal truncated fragment (without NLS domain, 97-261 aa) were also amplified and cloned into a bait vector as pGBKT7-TaHRC-N, pGBKT7-TaHRC-NΔNLS and pGBKT7-TaHRC-C. The co-transformaiton assays were performed as described by the manufacturer (Clontech Laboratories, CA, USA). All the primers used for vectors construction were listed in Supplemental Table S2.

#### Mutants generation by CRISPR/Cas9 gene editing

The single-guide RNA (sgRNA) was designed to target three different sites of *TaHRC* using the web-based E-CRISPR program (http://www.e-crisp.org/E-CRISP/). The details for construction of Cas9 and sgRNAs vectors as well as plant transformation, regeneration and selection processes for *TaHRC* mutants were described previously (Su et al., 2019). All primers used for vectors construction and mutants screening are listed in Supplemental Table S2.

#### FHB evaluation and statistical analysis

The detailed protocol of FHB evaluation for mutant lines was described previously (Su et al., 2019). In brief, the mutant plants were phenotyped for FHB resistance in a growth chamber by injecting a conidial spore suspension of *F. graminearum* (GZ3639) into a central spikelet in a spike at early anthesis using a syringe (Hamilton, Reno, NV). Ten spikes per line were inoculated in each replication and each experiment had three replications. The percentage of symptomatic spikelets (PSS) of FHB symptom in a spike was calculated at 14 days after inoculation. For box plots, boxes indicate the 25th–75th percentile, whiskers indicate the full data range, center lines indicate medians, crosses indicate means and the numbers inside boxes indicate sample size. *P-* values were generated from two-sided unpaired Student’s t-tests of the mean PSS of the mutant lines versus the mean PSS of the non-edited line.

#### Bimolecular fluorescence complementation (BiFC) assay

The full-length CDS of *TaHRC* and *TaCAXIP4* were amplified from ‘Clark’ and then cloned into the N-terminus and C-terminus of a split yellow fluorescent protein (YFP) in the expression vector pEarleygate201-YN and pEarleygate202-YC respectively, yielding YN-TaHRC and YC-TaCAXIP4 using the Gateway LR Clonase II enzyme mix kit (Invitrogen) as described previously (Lu et al., 2010). The mixed *Agrobacterium tumefaciens* (GV3101) cultures harboring equal concentrations of YN-TaHRC and YC-TaCAXIP4 vectors (final density OD600, 0.6) were infiltrated into 6-week-old epidermal *Nicotiana benthamiana* leaves. The signal of the reconstituted YFP fluorescence was observed and imaged at 48 hours after infiltration under a Zeiss LSM 880 confocal microscope (Carl Zeiss, Germany). All primers used for vectors construction are listed in Supplemental Table S2.

#### Subcellular colocalization assay

The full-length CDS of *TaHRC* and *TaCAXIP4* were cloned into in the expression vector pEarleygate103-GFP and pEarleygate102-CFP respectively, resulting in *TaHRC-GFP* and *TaCXIP4-CFP* using the Gateway LR Clonase II enzyme mix kit (Invitrogen) as described previously (Lu et al., 2010). The mixed *Agrobacterium tumefaciens* (GV3101) cultures harboring equal concentrations of *TaHRC-GFP* and *TaCXIP4-CFP* (final density OD600, 0.6) were infiltrated into 6-week-old epidermal *Nicotiana benthamiana* leaves. The GFP and CFP fluorescence signals at 48 hours after infiltration were observed and imaged under a Zeiss LSM 880 confocal microscope (Carl Zeiss, Germany) in two channels and merged using a lookup table with raw data in green and cyan colors, respectively. All primers used for vectors construction are listed in Supplemental Table S2.

#### Yeast Ca^2+^ supression assay

A Ca^2+^ sensitive yeast strain K667 (hypersensitive to high concentrations of Ca^2+^) was used for yeast transformation. The full-length CDS of *TaCAX1, TaCAXIP4* and *TaHRC* from ‘Clark’ were cloned into the yeast expression vector pGBKT7 and transformed/cotransformed into K667 yeast cells. Yeast cells expressing/coexpressing *TaCAX1*, *TaCAXIP4* and *TaHRC* were assayed on a yeast extract/peptone/dextrose (YPD) medium supplemented with and without 200 mM CaCl_2_, respectively. The detail protocol for yeast transformation was described previously (Cheng et al., 2004). All primers used for vectors construction are listed in Supplemental Table S2.

#### Reactive oxygen species (ROS) measurement

To examine whether *TaHRC* and *TaCaXIP4* affect ROS production in planta, ROS assay was performed in *N. benthamiana* leaves expressing *GFP* (Control), *TaHRC-GFP*, *TaCaXIP4-GFP* or both using a luminol-based chemiluminescence assay (Hao et al. 2019). *A. tumefaciens* GV3101 strains carrying *TaHRC-GFP* or *TaCaXIP4-GFP* were cultured overnight. The cultures were centrifuged, washed, and suspended in Agromix (10 mM MgCl_2_, 10 mM MES, and 100 μM acetosyringone). The cell suspension was infiltrated or co-infiltrated into 4- to 5-week-old *N. benthamiana* leaves with an OD600 of 0.4. Two days after infiltration, twelve leaf discs were removed from infiltrated zones with a cork borer and floated on water overnight. On the next day, water was replaced with 200 μL of the assay solution: 17 mM luminol L012, 1 μM horseradish peroxidase (Sigma, St. Louis, MO), and 100 μg/mL crab shell chitin (Sigma, St. Louis, MO). Luminescence was measured for 60 min using the Synergy HT and Gen5 software (BioTek Instruments, Inc. Winooski, VT).

---

**Table S1.**
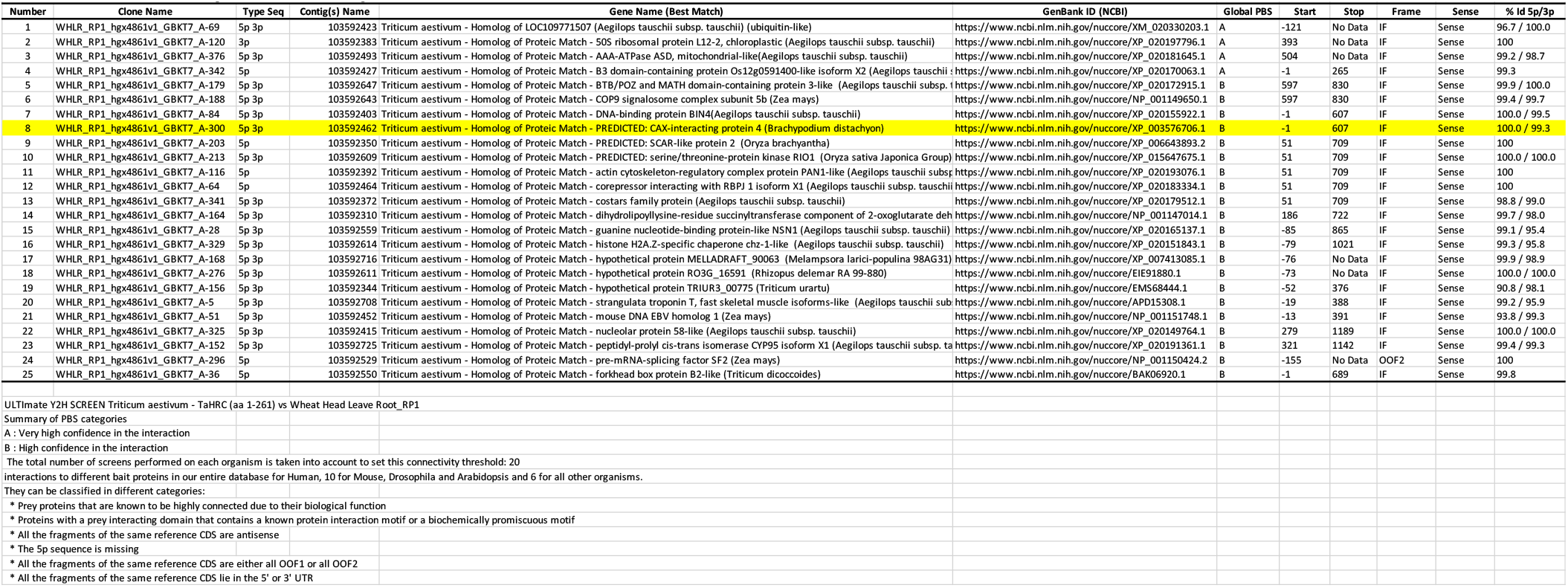
The results of ULTImate Y2H screen against wheat cDNA libraries using TaHRC as a bait.

**Table S2.**
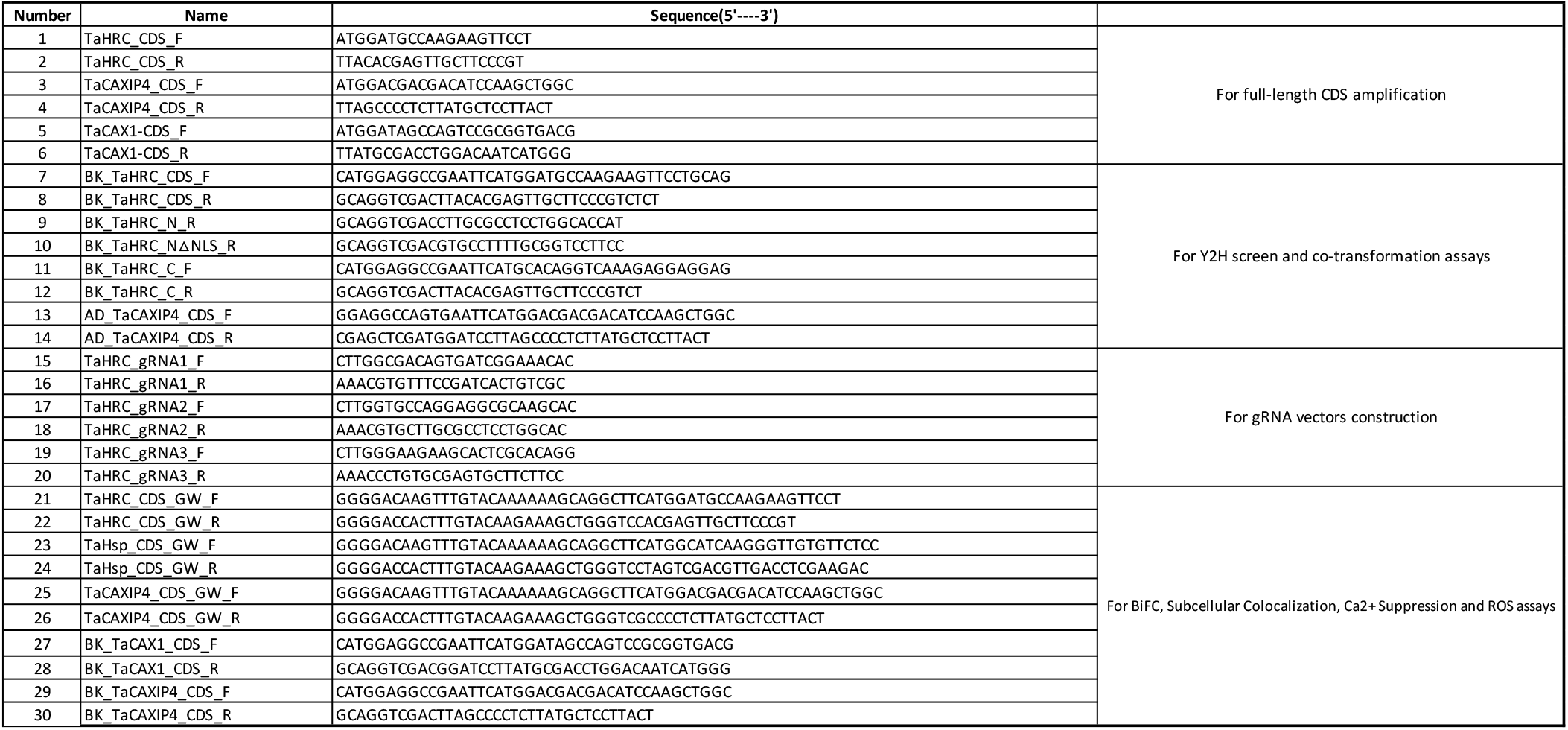
Primer sequences used in this study.

